# Reduced Resting Brain Entropy and Structural-Functional Reorganization in Hearing Loss

**DOI:** 10.64898/2026.11.16.766902

**Authors:** Ziliang Zhu, Zhen Cao, Donghui Song, Hengyi Rao

## Abstract

Hearing loss changes how the brain processes sound, yet how chronic auditory degradation shapes the temporal organization of spontaneous cortical activity remains unclear. We examined this question using multimodal MRI from 52 adults with age-related hearing loss and 30 normal-hearing controls. Guided by convergent neuroimaging evidence, we focused on the superior temporal gyrus (STG) and assessed brain entropy (BEN), complementary resting-state measures, and gray matter volume (GMV). Hearing loss was associated with lower BEN in a focal right-STG cluster after adjustment for local GMV (14 voxels; cluster FWER p = 0.008), consistent with more recurrent BOLD signal patterns. Six complementary measures of signal amplitude, local synchrony, symbolic sequence diversity, successive variation, and temporal persistence showed no corrected STG differences. A separate whole-brain analysis revealed higher bilateral midcingulate cortex (MCC) GMV (FWE p = 0.002 and 0.015), motivating examination of the auditory–cingulate relationship. The BEN-defined STG region showed stronger focal functional connectivity with MCC (FWE p = 0.031). Exploratory analyses suggested weaker directed STG-to-MCC influence alongside a small tractography-derived group difference. Exploratory directional comparisons indicated stronger right-MCC GMV–BEN coupling (p = 0.034) and STG– MCC SC–FC coupling (p = 0.037) in controls than in hearing loss. Together, these findings link greater temporal regularity in auditory association cortex to altered auditory–cingulate coordination and local structure– function correspondence, suggesting that adaptation to hearing loss involves both the organization of spontaneous cortical dynamics and their relationship to brain structure.

## Introduction

Hearing loss alters the sensory evidence available for speech and environmental sound. Reduced auditory fidelity changes activity within auditory cortex and increases the contribution of attention, prediction, and performance monitoring during listening. These adaptations extend beyond the auditory system: resting-state studies have reported cross-modal recruitment and altered large-scale network organization in hearing-impaired and deaf populations (Shang et al., 2020; Andin and Holmer, 2022), while the original analysis of the present cohort identified greater efficiency, density, and clustering in a visual subnetwork (Ponticorvo et al., 2022). A central question is therefore how chronic auditory degradation changes the intrinsic activity of auditory cortex before considering its wider network context.

The superior temporal gyrus (STG) is a compelling anatomical focus for investigating hearing-related cortical plasticity. Its role in encoding acoustic–phonetic features places it at a critical stage of speech processing (Mesgarani et al., 2014). Studies of hearing loss and deafness have documented altered local synchronization, recruitment of superior temporal regions by non-auditory inputs, and changes in their functional connectivity, suggesting that adaptation involves both local activity and interregional coordination (Li et al., 2013; Shiell et al., 2015; Andin and Holmer, 2022). Convergent evidence comes from a coordinate-based meta-analysis of 64 hearing-loss fMRI studies, encompassing 1,386 participants with hearing loss and 1,359 controls, which identified STG as the most consistent locus across activation, connectivity, and intrinsic-fluctuation studies (X. Wang et al., 2026). This convergence establishes a strong anatomical basis for examining STG and raises a more specific question about how hearing loss changes the temporal organization of activity within this region.

Temporal organization is central to the STG’s contribution to speech perception. Intracranial recordings show that STG responses encode acoustic–phonetic features and discrete landmarks in the speech envelope that help organize continuous speech into syllabic units (Mesgarani et al., 2014; Oganian and Chang, 2019). These representations also depend on the interaction between incoming sensory evidence and prior knowledge. During degraded-speech perception, prior knowledge modulates STG responses, with the timing of frontal and temporal effects supporting a top-down contribution to perceptual interpretation (Sohoglu et al., 2012). More recently, intracortical recordings demonstrated that semantic predictions align neuronal population dynamics supporting phonetic representations in anterior STG (Orepić et al., 2026). Together, these findings show that auditory representations depend on how cortical activity unfolds over time and how that unfolding is shaped by context. They motivate the hypothesis that prolonged degradation of auditory input may also alter the intrinsic temporal organization of STG activity. Resting-state measurements can address this question by characterizing the recurrence and predictability of spontaneous BOLD patterns, providing a complementary perspective on hearing-related plasticity beyond activity amplitude and interregional correlation.

Brain entropy (BEN) mapping estimates this temporal organization voxelwise using sample entropy (Wang et al., 2014). Sample entropy quantifies the conditional recurrence of short signal patterns (Richman and Moorman, 2000): lower BEN indicates that similar BOLD segments more often remain similar when extended, whereas higher BEN indicates less predictable pattern transitions. BEN therefore characterizes temporal irregularity under a defined sampling, filtering, and parameter regime. It has a reproducible cortical topography and high test-retest stability (Wang et al., 2014). Regional BEN also covaries with cerebral blood flow and low-frequency amplitude (Song et al., 2019a), relates to task-evoked activation and deactivation (Lin et al., 2022), and changes following focal neuromodulation (Song et al., 2019b, 2025). Together, these findings motivate a direct test of whether chronic auditory degradation alters the recurrence structure of spontaneous BOLD activity in STG.

Spontaneous BOLD activity has several complementary dimensions. ALFF and fALFF quantify low-frequency amplitude, ReHo measures local spatial concordance, Lempel-Ziv complexity characterizes diversity in a binarized symbol sequence, mean squared successive difference (MSSD) emphasizes moment-to-moment change, and intrinsic timescale measures temporal persistence (Zang et al., 2004, 2007; Zou et al., 2008; Wolff et al., 2022). Analyzing these measures together separates recurrence probability from oscillatory magnitude, neighborhood synchronization, symbolic diversity, successive variation, and autocorrelation. This multimetric design can reveal whether hearing loss changes a broad range of spontaneous signal properties or preferentially changes one aspect of STG temporal organization.

Local temporal dynamics arise within an anatomical and network context. Hearing loss is accompanied by regional gray matter changes, and local tissue variation can contribute to voxelwise BOLD measures. Separating BEN from co-localized morphology is therefore essential for identifying a temporal effect within STG. Once such an effect is established, whole-brain morphology and connectivity can reveal the wider structural and functional context in which it occurs.

We therefore tested whether hearing loss alters BOLD temporal regularity within left or right STG after accounting for local GMV, and whether complementary resting-state measures show corresponding changes. We separately examined whole-brain regional GMV. Any corrected STG BEN and whole-brain GMV findings were then carried forward to functional-connectivity analysis, followed by exploratory effective-connectivity, tractography, and cross-modal coupling analyses. We hypothesized that hearing loss would alter the recurrence structure of spontaneous activity within STG.

## Materials and Methods

### Participants and dataset

Data were obtained from the publicly available Hearing Loss Connectome dataset on OpenNeuro (ds005026, version 1.0.0; Ponticorvo and Esposito, 2024), originally described by Ponticorvo et al. (2022). The analyzed sample comprised 82 adults: 52 participants with age-related hearing loss (16 women; age, 63.67 ± 7.80 years) and 30 normal-hearing controls (20 women; age, 59.53 ± 7.17 years). Hearing-loss participants had received a clinical diagnosis within the preceding month and had not previously used hearing aids. Controls reported neither a history of hearing loss nor subjective hearing difficulties. Air-conduction thresholds were measured at 0.25, 0.5, 1, 2, 4, and 8 kHz, and the pure-tone average (PTA) was calculated separately for each ear across 0.5, 1, 2, and 4 kHz. PTA in the hearing-loss group was 50.3 ± 15.1 dB in the right ear and 50.3 ± 13.1 dB in the left ear. Pure-tone audiometry was available for 21 of the 30 controls, whose PTA was 15.2 ± 5.1 dB in the right ear and 14.5 ± 5.1 dB in the left ear. Age and PTA are expressed as mean ± SD. Speech audiometry was performed in 47 of the 52 hearing-loss participants (Ponticorvo et al., 2022). Participants had no reported neurological or psychiatric illness, prior ear surgery, or MRI contraindication. Ethical approval and written informed consent were obtained by the original investigators.

### MRI acquisition

MRI data were acquired on a 3 T Siemens MAGNETOM Skyra scanner with a 20-channel head/neck radiofrequency coil. The T1-weighted anatomical scan used a three-dimensional magnetization-prepared rapid gradient-echo sequence (TR = 2400 ms, TE = 2.25 ms, 1 mm isotropic voxels, 256 × 256 matrix, anterior-to-posterior phase encoding, and GRAPPA acceleration factor 2). Resting-state BOLD images were collected with multiband echo-planar imaging (TR = 1000 ms, TE = 30 ms, 2.5 mm isotropic voxels, 96 × 96 matrix, multiband factor 4, anterior-to-posterior phase encoding, 400 volumes, and total duration 6 min 40 s). Participants rested with their eyes open and were instructed to remain still. Diffusion MRI used multiband echo-planar imaging (TR = 4700 ms, TE = 91 ms, 2 mm isotropic voxels, 128 × 128 matrix, and multiband factor 2), with 64 diffusion directions at b = 1500 s/mm² and b = 0 images. Paired anterior-to-posterior and posterior-to-anterior reference acquisitions were collected for both modalities, comprising five BOLD volumes and five b = 0 diffusion volumes per phase-encoding direction, for susceptibility-distortion correction (Ponticorvo et al., 2022).

### Anatomical and functional preprocessing

Anatomical and resting-state functional images were preprocessed using fMRIPrep 23.1.4 (Esteban et al., 2019), built on Nipype 1.8.6 (Gorgolewski et al., 2011). T1-weighted images underwent intensity nonuniformity correction using N4BiasFieldCorrection (Tustison et al., 2010), brain extraction using ANTs, and tissue segmentation into gray matter, white matter, and cerebrospinal fluid using FSL FAST (Zhang et al., 2001; Jenkinson et al., 2012). Nonlinear registration to the MNI152NLin6Asym and MNI152NLin2009cAsym templates was performed using ANTs (Avants et al., 2011).

For each resting-state run, a BOLD reference image was generated, head motion was estimated using FSL MCFLIRT (Jenkinson et al., 2002), and slice timing was corrected using AFNI 3dTshift (Cox, 1996). The BOLD reference was coregistered to the T1-weighted image using FreeSurfer mri_coreg followed by FSL boundary-based registration (Greve and Fischl, 2009). Motion, coregistration, and spatial-normalization transforms were composed to minimize interpolation, and volumetric resampling used Lanczos interpolation. Subsequent functional analyses used images in MNI152NLin2009cAsym space.

Additional nuisance processing was performed using Nilearn (Abraham et al., 2014). Regressors included white-matter and cerebrospinal-fluid signals and their temporal derivatives, standardized DVARS, framewise displacement, root-mean-square displacement, the first three CSF-derived anatomical CompCor components (Behzadi et al., 2007), an additional nuisance regressor labeled “tcompcor” in the supplied confounds files, and the six rigid-body motion parameters and their temporal derivatives. For BEN and FC, BOLD time series were linearly detrended, standardized, and band-pass filtered at 0.009–0.08 Hz. For BEN estimation, the processed BOLD images were smoothed using a 6-mm full-width-at-half-maximum Gaussian kernel. The first four volumes were excluded before BEN estimation and functional-connectivity calculation, leaving 396 time points. Mean framewise displacement was calculated over these retained volumes and included as a covariate in subsequent group-level functional analyses.

### Gray matter volume

Gray matter volume (GMV) was estimated using a combined fMRIPrep–FSL-VBM workflow, following our previous study (Song and Wang, 2026). This approach integrates fMRIPrep-based tissue segmentation with FSL-based spatial registration. A systematic comparison identified fMRIPrep–FSL and CAT12.8 as effective pipelines for capturing age-related variation in regional GMV and predicting age (Antonopoulos et al., 2023).

Gray-matter probability maps obtained from anatomical preprocessing were processed using the optimized FSL-VBM protocol (Good et al., 2001; Douaud et al., 2007). The maps were initially affine registered to the ICBM-152 gray-matter template and used to construct a left–right symmetric study-specific template. Individual gray-matter maps were then nonlinearly registered to this template using FNIRT and modulated using the Jacobian determinants of the nonlinear transformations to preserve local tissue volume. The modulated maps were smoothed with an 8-mm full-width-at-half-maximum Gaussian kernel. Total intracranial volume (TIV) was calculated by summing the native-space gray-matter, white-matter, and cerebrospinal-fluid probability maps and multiplying by voxel volume.

For whole-brain regional analysis, mean modulated GMV was extracted from the 116 regions of the Automated Anatomical Labeling (AAL) atlas (Tzourio-Mazoyer et al., 2002). Group differences were assessed using linear models with hearing-loss group as the predictor of interest and age, sex, and TIV as covariates. Two-sided family-wise error correction across all 116 regions was performed using 10,000 Freedman–Lane permutations, with the maximum absolute t statistic across regions recorded in each permutation (Winkler et al., 2014).

Supportive voxelwise GMV analyses used models adjusting for age, sex, and TIV. Clusters were formed at two-sided voxelwise p < 0.001 using 26-neighbor connectivity. Maximum-cluster-extent FWER correction used 5,000 nuisance-residual permutations, separately within the bilateral AAL MCC mask and the whole-brain mask. The MCC-restricted analysis characterized the localization of the whole-parcellation finding.

### Brain entropy mapping

Voxelwise BEN maps were calculated from the nuisance-processed, 6-mm-smoothed resting-state BOLD time series using sample entropy (Richman and Moorman, 2000; Wang et al., 2014). The embedding dimension was set to m = 3, and the tolerance was set to r = 0.60 times the standard deviation of each voxel’s time series. The first four volumes were excluded, leaving 396 observations for BEN estimation. Individual BEN maps were resampled to a 3-mm isotropic MNI152 grid and smoothed with an 8-mm full-width-at-half-maximum Gaussian kernel.

Group differences in BEN were assessed using voxelwise general linear models, with hearing-loss group as the predictor of interest and age, sex, mean framewise displacement over the retained volumes, TIV, and co-localized GMV as covariates. The left and right STG were defined using the AAL atlas and analyzed separately (Tzourio-Mazoyer et al., 2002). Within each STG mask, clusters were identified using a two-sided voxelwise cluster-forming threshold of p < 0.001 and 26-neighbor connectivity. Cluster-level family-wise error was controlled using 5,000 nuisance-residual permutations, with the maximum cluster extent within the mask recorded in each permutation (Nichols and Holmes, 2002; Winkler et al., 2014). Clusters with FWER-corrected p < 0.05 were considered significant.

### Alternative resting-state measures

Six complementary resting-state measures were calculated to characterize signal amplitude, local synchrony, symbolic sequence diversity, successive variation, and temporal persistence: amplitude of low-frequency fluctuations (ALFF), fractional ALFF (fALFF), regional homogeneity (ReHo), Lempel–Ziv complexity (LZC), mean squared successive difference (MSSD), and intrinsic timescale (ITS). Calculations used the spatially normalized BOLD images after exclusion of the first four volumes, leaving 396 observations. The retained time series underwent linear detrending and regression of the nuisance variables described above.

ALFF was calculated as the mean Fourier amplitude within 0.009–0.08 Hz (Zang et al., 2007). fALFF was calculated as the summed amplitude within this band divided by the summed amplitude across nonzero frequencies up to 0.25 Hz (Zou et al., 2008). Both measures were derived from the nuisance-regressed time series before band-pass filtering. For ReHo, the time series were filtered at 0.009–0.08 Hz using a fourth-order Butterworth filter applied in forward and reverse directions. ReHo was then calculated using Kendall’s coefficient of concordance across each voxel and its 26 immediate neighbors, forming a 27-voxel neighborhood (Zang et al., 2004).

LZC, MSSD, and ITS were calculated from nuisance-regressed, temporally standardized time series without additional band-pass filtering. For LZC, each voxel’s time series was binarized using its temporal median, and sequence diversity was quantified using Lempel–Ziv 1976 parsing (Lempel and Ziv, 1976). The resulting phrase count was normalized as c(n)log (n)/n, where c(n) is the phrase count and n are the number of observations. LZC was calculated within the bilateral AAL STG masks and a surrounding three-voxel margin to support subsequent spatial smoothing. MSSD was calculated as the mean squared difference between consecutive standardized observations, quantifying successive signal variation (Garrett et al., 2013). ITS was estimated from the initial positive portion of the autocorrelation function, following the autocorrelation-area approach of Watanabe et al. (2019). Specifically, autocorrelations were summed from lag zero to the lag preceding the first nonpositive value, with a maximum lag of 50 s, and multiplied by the repetition time to express ITS in seconds.

Individual ALFF, fALFF, ReHo, MSSD, and ITS maps were spatially standardized within their valid brain masks and smoothed with an 8-mm full-width-at-half-maximum Gaussian kernel. Normalized LZC maps were smoothed using the same kernel, with support-weighted normalization at mask boundaries. Analyses were restricted to voxels with valid metric coverage in at least 80% of participants.

Voxelwise group differences were assessed separately within the left and right AAL STG masks, adjusting for age, sex, mean framewise displacement, TIV, and co-localized GMV. As in the BEN analysis, clusters were formed at a two-sided voxelwise threshold of p < 0.001 using 26-neighbor connectivity, and cluster-level FWER was controlled within each STG mask using 5,000 nuisance-residual maximum-cluster-extent permutations (Winkler et al., 2014). Complementary whole-brain regional analyses were performed for ALFF, fALFF, ReHo, MSSD, and ITS across 116 AAL regions, adjusting for the same participant-level covariates and regional GMV. Joint FWER correction across the five measures and 116 regions (580 tests) used 10,000 permutations and the maximum absolute t statistic across all tests.

### Seed-based functional connectivity

Seed-based FC analysis was performed to characterize the connectivity of the BEN-defined right STG region (Biswal et al., 1995). The seed comprised the 14-voxel cluster showing lower BEN in the hearing-loss group after adjustment for demographic covariates, head motion, TIV, and co-localized GMV, which survived cluster-level FWER correction within the right AAL STG mask. This binary cluster mask was resampled to each participant’s functional-image grid using nearest-neighbor interpolation and intersected with the participant’s brain mask. The mean seed time series was extracted from the nuisance-processed BOLD images after removal of the first four volumes. Pearson correlations were calculated between the seed time series and the time series of each valid brain voxel, yielding individual whole-brain FC maps. Correlation coefficients were Fisher-z transformed, and the resulting maps were smoothed with an 8-mm full-width-at-half-maximum Gaussian kernel.

Group differences were assessed using voxelwise general linear models with hearing-loss group as the predictor of interest and age, sex, mean framewise displacement over the retained volumes, and TIV as covariates. The analysis mask included voxels with valid FC estimates in at least 80% of participants, excluding the seed and its immediate neighborhood. The bilateral MCC was selected as the anatomical search region based on the whole-brain AAL regional GMV findings and defined as the union of the left and right AAL middle cingulate regions (Tzourio-Mazoyer et al., 2002). Within this combined mask, two-sided voxelwise FWE correction was performed using 10,000 Freedman–Lane nuisance-residual permutations. The maximum absolute t statistic across all voxels in the bilateral MCC mask was recorded in each permutation to construct the null distribution (Nichols and Holmes, 2002; Winkler et al., 2014). Voxels with FWE-corrected p < 0.05 were considered significant. Complementary whole-brain inference used two-sided voxelwise Bonferroni correction across the analysis mask.

A leave-one-out sensitivity analysis evaluated the stability of MCC target selection. In each fold, the five MCC voxels with the largest positive group-effect t statistics were selected using the other 81 participants, and their mean FC was extracted from the held-out participant. These held-out values were analyzed using the same group-level covariates. The BEN-defined STG seed remained fixed across folds.

### Spectral dynamic causal modeling

Exploratory effective-connectivity analyses used spectral DCM in SPM25 (25.01.02) with MATLAB R2021a, fitting a generative model of neuronal and hemodynamic dynamics to regional BOLD cross-spectral densities (Friston et al., 2014).

Two separate two-node models were constructed, pairing the right STG with either the left or right MCC. The STG node was defined by the 14-voxel BEN cluster used in the FC analysis. The MCC nodes comprised the complete left and right middle cingulate regions of the AAL atlas, respectively (Tzourio-Mazoyer et al., 2002). ROI masks were resampled to the functional-image grid using nearest-neighbor interpolation. For each participant, the mean BOLD time series was extracted from each ROI after exclusion of the first four volumes, yielding 396 observations. Regional time series underwent nuisance regression using the confounds described above, linear detrending, and temporal standardization, without additional band-pass filtering.

Each model was fully connected, including reciprocal STG–MCC connections and a self-connection for each node. A linear, one-state neuronal model was specified, with no exogenous inputs or modulatory effects. Subject-level models were estimated using the cross-spectral density inversion implemented in SPM.

Group effects on the endogenous connectivity parameters were assessed separately for the left- and right-MCC models using parametric empirical Bayes (PEB). The second-level design matrix included an intercept, mean-centered group membership (hearing loss = 1; normal hearing = 0), standardized age, and mean-centered sex. Bayesian model reduction and Bayesian model averaging were then applied to obtain posterior estimates of the group effects (Zeidman et al., 2019). Posterior probabilities ≥ 0.95 were used as the criterion for strong evidence. Model fit was summarized by the proportion of explained spectral variance, and the PEB analysis was repeated after excluding models with explained variance below 70%.

A complementary analysis paired the same right-STG node with the five-voxel MCC cluster surviving voxelwise FWE correction in the FC analysis. Time-series extraction, model specification, and group-level estimation followed the same procedures.

### Probabilistic tractography

Diffusion images were preprocessed with QSIPrep 26.0.0 at 2-mm output resolution, and ACPC-aligned diffusion images, gradient directions, and brain masks were used for fiber-orientation estimation with FSL bedpostx_gpu (Cieslak et al., 2021; Behrens et al., 2007). The 14-voxel right-STG BEN seed was transformed into each participant’s diffusion space using the QSIPrep-derived MNI-to-ACPC transform with nearest-neighbor interpolation and intersected with the diffusion brain mask. Whole-brain tracking from the seed used probtrackx2_gpu with 5,000 samples per seed voxel, 2,000 maximum steps, a 0.5-mm step length, curvature threshold 0.2, subsidiary fiber-volume threshold 0.01, loop checking, and distance correction. Tract-distribution maps were divided by waytotal, the number of accepted samples, and transformed to the standard-space FC grid using linear interpolation. The target was the 28-voxel exploratory MCC cluster from the earlier FC map (uncorrected voxelwise p < 0.001; minimum cluster extent 10 voxels). Mean and maximum normalized tract values and the fraction of target voxels with nonzero values were extracted. Group models adjusted for age, sex, and diffusion-space STG seed voxel count, with HC3 robust standard errors and two-sided t tests (MacKinnon and White, 1985).

### Multimodal structure-function coupling

Structure–function coupling was examined across participants at local and interregional levels. Local analyses quantified the relationship between regional GMV and BEN, whereas interregional analyses assessed the relationship between SC and FC along the STG–MCC pathway.

Local GMV–BEN coupling was evaluated within the 14-voxel right-STG BEN cluster and the AAL-defined right MCC, with bilateral MCC examined as a complementary regional summary. GMV and BEN maps each underwent 8-mm FWHM map-level smoothing. GMV maps were resampled to the 3-mm BEN grid using continuous interpolation, and mean GMV and BEN were extracted from identical regional masks. Within each group, both variables were separately residualized for age, sex, and mean framewise displacement. Pearson correlations between the residuals provided group-specific partial correlation coefficients.

Interregional SC–FC coupling was evaluated using the right-STG seed and the 28-voxel exploratory MCC target. SC was quantified as the mean waytotal-normalized tract-distribution value within the MCC target from the final QSIPrep-based tractography analysis. Values were transformed as ln(1 + 10 × SC). FC was quantified as the mean Fisher-z-transformed seed-to-voxel correlation within the same target. Group-specific partial correlations controlled for age, sex, mean framewise displacement, TIV, and the number of STG seed voxels in diffusion space.

Within-group partial correlations were evaluated using two-sided t tests, with t = r√[(n − k − 2)/(1 − r²)] and n − k − 2 degrees of freedom, where n denotes group sample size and k denotes the number of covariates. Differences between group-specific partial correlations were assessed using an approximate Fisher r-to-z comparison (Fisher, 1921). The difference between Fisher-transformed correlations was divided by √[1/(nNH − k − 3) + 1/(nHL − k − 3)], with k = 3 for local GMV–BEN coupling and k = 5 for interregional SC–FC coupling. Exploratory one-sided comparisons evaluated whether coupling was stronger in normal-hearing controls than in participants with hearing loss. Coupling p values were reported without multiplicity correction.

Sensitivity of right-MCC GMV–BEN coupling to unequal group sizes was examined using 10,000 balanced subsamples. Each iteration retained all 30 controls and randomly selected 30 of the 52 hearing-loss participants without replacement. Covariate adjustment and partial correlations were recalculated within each subsample. The between-group correlation difference was summarized by its median, 2.5th–97.5th percentile range, and proportion of iterations showing stronger coupling in controls.

### Robustness analyses

BEN seed stability was assessed using leave-one-out selection within right AAL STG. For each held-out participant, the voxelwise group model was fitted to the remaining 81 participants with age, sex, mean FD, TIV, and voxelwise GMV adjustment. Negative clusters were selected at two-sided voxelwise p < 0.001, 26-neighbor connectivity, and minimum extent 10 voxels. BEN and co-localized GMV were extracted for the held-out participant; folds without a qualifying cluster were omitted from the held-out summary model. Voxel-selection frequency used all 82 folds as its denominator. Motion sensitivity of BEN and FC summaries was assessed by adding squared mean FD or excluding participants above the 95th or 90th percentile of mean FD. BEN summary models included age, sex, mean FD, TIV, and local GMV; FC summary models included age, sex, mean FD, and TIV. The FC leave-one-out procedure is described above. Summary statistics are reported in Supplementary Table S5.

## Results

### Focal BEN reduction in the right STG

Figure 1 summarizes the corrected right-STG BEN, bilateral-MCC GMV, and focal STG–MCC FC findings, together with participant-level distributions and supporting leave-one-out analyses.

**Figure 1.**
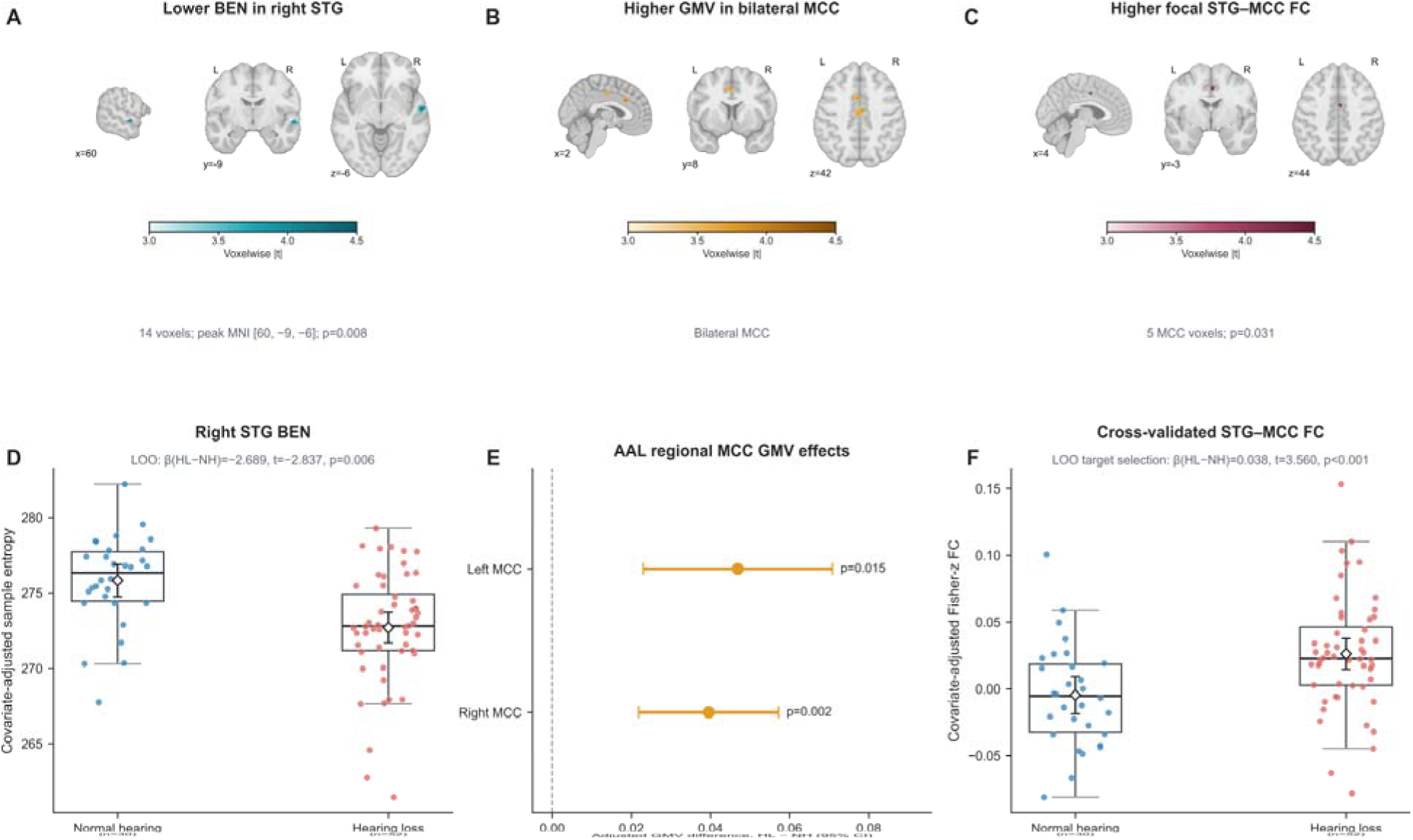
Primary right superior temporal gyrus BEN finding and data-driven midcingulate results. (A) A 14-voxel right superior temporal gyrus (STG) cluster showed lower BOLD time-series entropy (BEN) in participants with hearing loss (HL) than in participants with normal hearing (NH). The model included age, sex mean framewise displacement (FD), total intracranial volume (TIV), and voxelwise 8-mm local gray matter volume (GMV); the cluster survived right-STG maximum-cluster permutation family-wise error rate correction (FWER p=0.008; peak MNI coordinate [60, −9, −6]). BEN indexes the temporal irregularity of the BOLD signal under the stated preprocessing and parameter settings. (B) Modulated 8-mm GMV was higher in bilateral midcingulate cortex (MCC). The displayed clusters survived maximum-cluster FWER correction within the bilateral AAL MCC mask. In the complementary whole-brain AAL regional analysis, right and left MCC survived maxT correction across 116 regions (right: beta=0.0395, t=4.451, p_FWE=0.002; left: beta=0.0469, t=3.908, p_FWE=0.015). (C) Fisher-z-transformed functional connectivity (FC) from the right-STG BEN cluster was higher in a focal five-voxel MCC cluster, surviving permutation maximum-|t| FWER correction across the bilateral MCC (peak t=4.247, p_FWE=0.031). Colorbars in A-C show voxelwise absolute group-effect t values on the same 3.0-4.5 scale; A depicts HL < NH, whereas B and C depict HL > NH. (D) Full-sample seed BEN values after removal of nuisance variance; the LOO statistic summarizes a separate held-out analysis. Leave-one-out (LOO) cluster selection retained the group effect (beta_HL-NH=-2.69, t=-2.84, p=0.006; 79 evaluable folds); 12 of the 14 full-sample cluster voxels were selected in at least 80% of folds. (E) AAL regional MCC GMV effects with 95% confidence intervals. (F) Participant-level STG-MCC FC after LOO selection of the five MCC target voxels using the other 81 participants (beta_HL-NH=0.0376, t=3.56, p<0.001). Points represent participants; diamonds and bars show group means and approximate 95% confidence intervals. Participant distributions are descriptive; corrected spatial inference is reported in panels A-C. NH n=30; HL n=52. Boxes show the interquartile range and median; whiskers extend to observations within 1.5 times the interquartile range.

Hearing loss was associated with lower BEN in a focal right STG cluster after voxelwise adjustment for 8-mm GMV. A 14-voxel cluster exceeded the voxelwise cluster-forming threshold of p < 0.001 and survived right-STG maximum-extent permutation correction (FWER p = 0.008). The effect was confined to the right STG and was diluted when signal was averaged across broader anatomical STG regions.

### Specificity across complementary resting-state measures

None of the six complementary measures produced a corrected cluster in either STG with 8-mm voxelwise GMV adjustment. The uncorrected candidates included ALFF (R STG, 6 voxels; peak t = 3.790, cluster FWER p = 0.060); Lempel-Ziv complexity (L STG, 2 voxels; peak t = 3.700, cluster FWER p = 0.139); intrinsic timescale (L STG, 1 voxel; peak t = 3.429, cluster FWER p = 0.105). Full STG results are provided in Supplementary Table S1.

The whole-brain AAL analysis likewise yielded no effect after joint correction across 580 metric-by-region tests. The lowest corrected p values occurred for lower intrinsic timescale in left posterior cingulate cortex (t = −3.850, p_FWE = 0.089), higher ReHo in left cerebellar lobule VIII (t = 3.832, p_FWE = 0.095), and higher MSSD in left posterior cingulate cortex (t = 3.812, p_FWE = 0.101). Metric-wise FDR sensitivity analyses identified 10 regional effects after 8-mm GMV adjustment, predominantly involving cerebellar ReHo. The complete regional estimates are reported in Supplementary Table S3. These effects did not survive the joint five-metric correction. The strongest result for each metric is summarized in Supplementary Table S2 and Supplementary Fig. S1; all 580 regional tests are provided in Supplementary Data S1.

### Higher bilateral MCC gray matter volume

The whole-parcellation analysis of 8-mm GMV identified higher GMV in right MCC (beta = 0.0395, t = 4.451, FWE p = 0.002) and left MCC (beta = 0.0469, t = 3.908, FWE p = 0.015) after max-|t| correction across all 116 AAL regions. MCC-restricted voxelwise analyses localized the regional effect and were used as supportive anatomical characterization. Regional estimates are shown in Fig. 1E; supportive voxelwise localization is shown in Fig. 1B.

### Stronger focal right STG-MCC functional connectivity

Seed-based analysis of the right STG BEN cluster revealed stronger FC in hearing loss within a focal MCC locus. Five MCC voxels survived permutation max-|t| correction within the bilateral MCC mask. The peak was located near MNI coordinates x = 3.5, y = −2.5, z = 44.0 (t = 4.247; FWE p = 0.031). Averaging across the full anatomical MCC attenuated the effect (beta = 0.0060, t = 1.260, permutation p = 0.211), consistent with a spatially focused change in STG-MCC coordination. The focal result is shown in Fig. 1C, and held-out FC values are shown in Fig. 1F.

### Effective and structural connectivity

Figure 2 summarizes robustness analyses and the multimodal characterization of the STG-MCC association.

**Figure 2.**
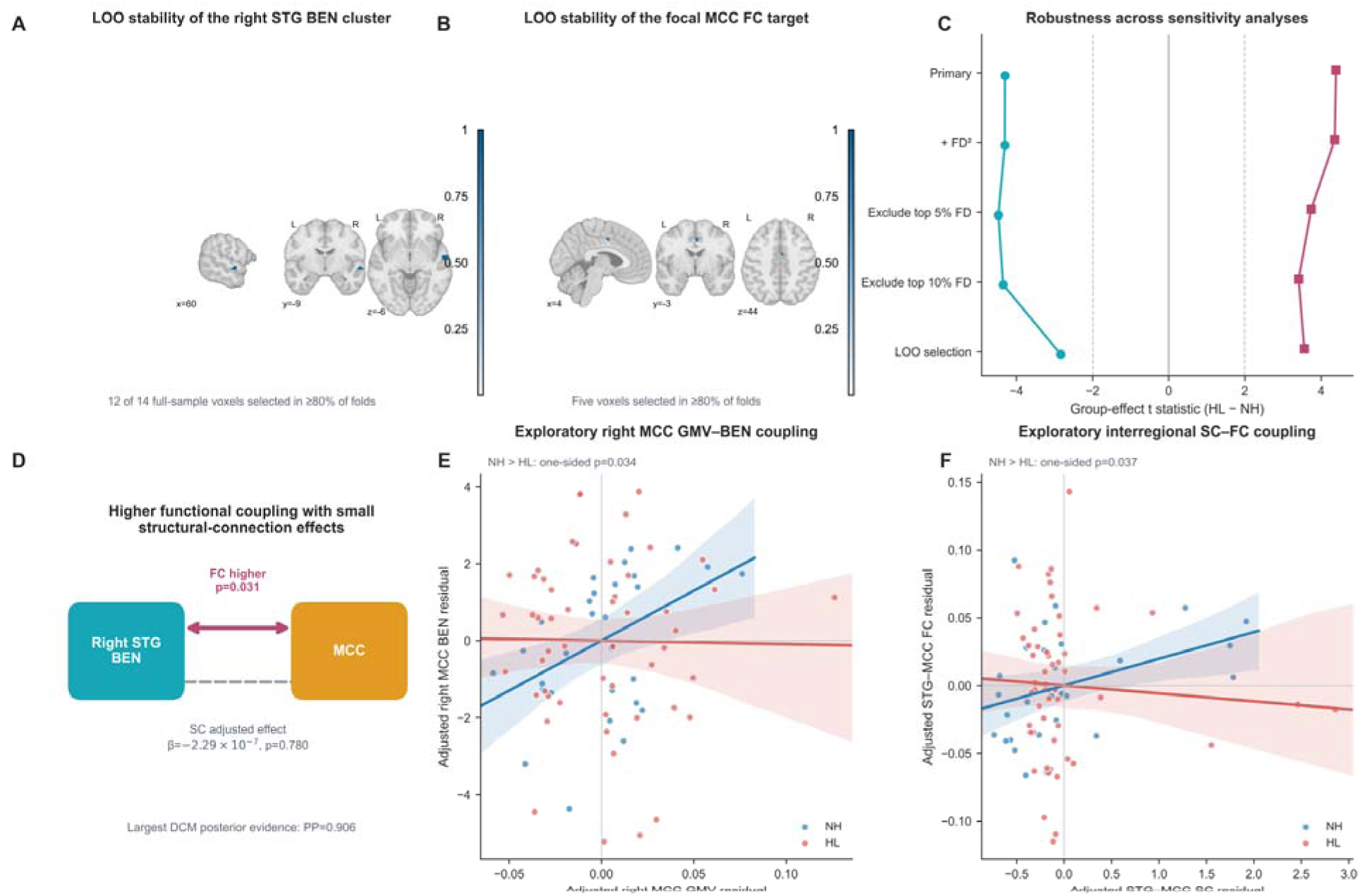
Robustness and multimodal characterization of the data-driven STG-MCC association. (A,B) Voxel selection frequency across 82 LOO folds (79 evaluable BEN folds) for the right-STG BEN cluster and focal MCC FC target. (C) Group-effect t statistics under the primary model, an additional quadratic-FD model, exclusion of participants in the highest 5% or 10% of FD, and LOO selection. Dashed vertical lines at approximately |t|=1.99 provide a visual reference and do not represent a multiple-comparison threshold. (D) Summary of multimodal connectivity evidence. Focal undirected STG-MCC FC was higher in HL. The age-, sex-, and seed-size-adjusted QSIPrep tractography effect was small (beta=-2.29 x 10^-7, p=0.780), and the largest posterior probability among the displayed spectral DCM parameters was 0.906. (E) Covariate-adjusted right-MCC GMV-BEN associations from independently generated 8-mm maps (NH: partial r=0.418, p=0.030; HL: partial r=-0.014, p=0.924; directional NH > HL Fisher comparison: z=1.824, one-sided p=0.034). (F) Covariate-adjusted interregional SC-FC associations (NH: partial r=0.377, p=0.063; HL: partial r=-0.070, p=0.642; directional NH > HL Fisher comparison: z=1.787, one-sided p=0.037). Lines and shaded bands in panels E and F show group-specific fits and 95% confidence bands.

All 82 models converged in both atlas-based DCM analyses, with median explained spectral variance of 91.4% for the left-MCC model and 91.6% for the right-MCC model. The estimated group difference was negative for right STG-to-right MCC effective connectivity (estimate = −0.305, posterior probability = 0.884) and remained similar after excluding the participant with explained variance below 70% (estimate = −0.317, posterior probability = 0.895). The corresponding left-hemisphere estimate was smaller (estimate = −0.171, posterior probability = 0.685). The right-sided estimate therefore provided directional context for the FC result, with evidence below the 0.95 criterion. In the post hoc focal-MCC model, neither between-region direction showed a group effect; its largest posterior probability was 0.906 for increased MCC self-inhibition after exclusion of the low-fit participant. Model estimates and posterior probabilities are summarized in Supplementary Fig. S2.

Probabilistic tractography of the exploratory 28-voxel MCC target showed no group difference. The adjusted hearing-loss effect was small for mean tract probability (beta = −2.29 x 10^-7, t = −0.281, p = 0.780), MCC nonzero fraction (beta = 0.00104, t = 0.184, p = 0.855), and maximum tract probability (beta = −1.45 x 10^-5, t = −0.848, p = 0.399). These exploratory tract measures are summarized in Supplementary Fig. S3.

### Structure-function coupling

Local and interregional analyses showed a convergent pattern of stronger structure–function coupling in normal-hearing controls. Within right MCC, GMV and BEN were positively correlated in controls after adjustment for age, sex, and head motion (partial r = 0.418, df = 25, p = 0.030), whereas the association was near zero in hearing loss (partial r = −0.014, df = 47, p = 0.924). The exploratory directional Fisher comparison indicated stronger coupling in controls (z = 1.824, one-sided p = 0.034; Fig. 2E). This ordering persisted across all 10,000 balanced subsamples, with a median correlation difference of 0.435 and a 2.5th–97.5th percentile range of 0.234–0.667.

The complementary bilateral-MCC analysis showed the same pattern: GMV–BEN coupling was positive in controls (partial r = 0.388, p = 0.046) and weakly negative in hearing loss (partial r = −0.057, p = 0.697), with a stronger correlation in controls in the directional comparison (z = 1.850, one-sided p = 0.032). Within the right-STG BEN cluster, associations were weak in both groups (controls: partial r = 0.009, p = 0.965; hearing loss: partial r = 0.115, p = 0.430); the NH > HL comparison yielded z = −0.425 and one-sided p = 0.664 (Supplementary Table S4).

Interregional coupling based on the final QSIPrep tractography results was positive in controls (partial r = 0.377, df = 23, p = 0.063) and weakly negative in hearing loss (partial r = −0.070, df = 45, p = 0.642). The exploratory directional Fisher comparison indicated stronger SC–FC coupling in controls (z = 1.787, one-sided p = 0.037; Fig. 2F). Together, the local MCC and interregional findings suggest reduced correspondence between structural variation and spontaneous functional properties in hearing loss (Supplementary Fig. S4).

## Discussion

This study identifies a focal shift toward more recurrent spontaneous dynamics in right auditory association cortex and situates it within a data-driven STG-MCC reorganization. Lower BEN persisted after local GMV adjustment and was not reproduced by measures of amplitude, local synchrony, symbolic diversity, successive variation, or temporal persistence, identifying pattern recurrence as the clearest corrected STG finding in the multimetric assessment. Independent whole-brain analysis identified higher bilateral MCC GMV, and the BEN-defined STG locus showed stronger focal connectivity with MCC. Together, these findings connect altered auditory-cortical temporal organization to cingulate morphology and coordination identified through a separate whole-brain regional analysis.

The right-STG BEN reduction adds a temporal-statistical dimension to established accounts of auditory-cortical reorganization. Previous studies have reported altered synchronization within superior temporal cortex, abnormal low-frequency amplitude in temporal and association regions, and changes in neurochemical balance and dynamic network states in hearing loss (Li et al., 2013; Yang et al., 2014; Li et al., 2023; Su et al., 2024). The recent meta-analysis by Wang et al. (2026) strengthens the anatomical context for the present finding by identifying STG as the most reproducible locus across hearing-loss activation, connectivity, and intrinsic-fluctuation studies. Within this region, lower BEN indicates that short BOLD patterns were more likely to recur in hearing loss.

Predictive processing provides a mechanistic framework for this temporal result. When prior knowledge improves the clarity of degraded speech, frontal signals precede changes in STG activity; perceptual learning produces convergent reductions in STG responses, showing that immediate expectations and longer-term experience reshape auditory representations through a shared cortical mechanism (Sohoglu et al., 2012; Sohoglu and Davis, 2016). At the neuronal-population level, Orepić et al. (2026) further showed that semantic predictions align ongoing phonetic dynamics in human anterior STG. Together, these studies position STG as an interface where bottom-up sensory evidence is continuously reconciled with internally generated predictions. Chronic loss of precise auditory input may reorganize this balance and stabilize a more recurrent set of spontaneous configurations, expressed here as lower resting-state BEN. This account predicts that individual differences in STG BEN will track contextual benefit and speech-in-noise performance, linking intrinsic temporal organization to predictive listening under degraded input.

The multimetric panel sharpens the interpretation of the STG effect. ALFF and fALFF index low-frequency amplitude, ReHo indexes neighborhood concordance, Lempel-Ziv complexity summarizes symbolic sequence diversity, MSSD emphasizes successive change, and intrinsic timescale captures autocorrelation persistence. The localization of the corrected group difference to BEN places particular emphasis on altered BOLD pattern recurrence in this cohort. This dissociation gives the BEN result a specific temporal meaning rather than treating it as a generic marker of abnormal spontaneous activity. Whole-brain sensitivity analyses identified their largest effects in posterior cingulate, cerebellar, and superior-frontal regions; these secondary patterns are reported in the Supplementary Materials alongside the corrected STG findings.

The bilateral MCC GMV increase emerged independently at the whole-brain regional level. Structural studies of hearing loss have often emphasized lower GMV in auditory cortex, posterior cingulate cortex, and precuneus, although increases in non-auditory association regions have also been reported (Yang et al., 2014; Wang et al., 2016; Alfandari et al., 2018). The present result adds to this regionally heterogeneous pattern rather than implying a uniform direction of structural change. MCC participates in performance monitoring and action selection under uncertainty (Vogt, 2016). During difficult speech recognition, cingulo-opercular activity increases after errors and predicts better recognition on the next trial, including in older adults (Vaden et al., 2015). Sustained recruitment of these control processes offers a plausible functional setting for the MCC finding, although the cross-sectional GMV effect cannot establish whether it reflects adaptation to hearing loss or a pre-existing difference.

The focal increase in right STG-MCC FC provides functional evidence for an auditory-cingulate association in the same cohort. Fitzhugh et al. (2019) reported stronger connectivity between right Heschl’s gyrus and dorsal anterior cingulate cortex as hearing declined, even after accounting for age and cognitive variation. Deaf adults also show stronger connectivity between temporal cortex and cingulate or insular regions (Andin and Holmer, 2022). The present analysis extends this literature by using a temporally altered STG locus as the seed and an separately identified structural MCC finding as the anatomical search space. At a broader scale, the original analysis of this cohort identified greater efficiency, density, and clustering in a visual subnetwork (Ponticorvo et al., 2022), while another hearing-loss cohort showed contraction of the principal cortical functional gradient (Tong et al., 2023). The focal STG-MCC effect may represent one local expression of this wider redistribution of communication between sensory and control systems.

FC, DCM, and tractography characterize different properties of the same regional pair. Stronger FC indicates greater temporal covariation between STG and MCC but does not specify the direction or anatomical route of that covariation. Spectral DCM estimated a weaker right STG-to-MCC influence, with posterior evidence below the prespecified threshold, whereas probabilistic tractography showed no detectable group difference in pathway strength. The pattern is compatible with altered functional use of a broadly similar macroscopic structural scaffold. Greater zero-lag coordination could coexist with weaker unique STG-to-MCC influence if the regions receive more consistent common input or participate more often in the same distributed state. The present data do not identify which of these mechanisms produced the FC increase.

The coupling analyses extend the regional findings by showing that hearing loss is associated with changes in how structural variation relates to spontaneous functional activity. Recent work in presbycusis has linked regional ALFF-to-GMV ratios to hearing thresholds and cognitive performance, highlighting the relevance of local structure–function relationships to hearing-related adaptation (Li et al., 2026). Our analysis extends this perspective to the correspondence between regional morphology and BOLD pattern recurrence. Within right MCC, greater GMV was associated with higher BEN in normal-hearing controls, whereas this relationship was near zero in hearing loss. The exploratory directional comparison supported stronger coupling in controls, and the same ordering persisted across all balanced subsamples. This finding adds a relational dimension to the higher MCC GMV observed in hearing loss: regional morphology and BOLD pattern recurrence showed different correspondence between groups. Previous work has identified predominantly negative associations of BEN with GMV and surface area across several cortical territories (Del Mauro and Wang, 2025). The positive association observed here in control MCC emphasizes that morphology–entropy relationships depend on regional organization and the population examined. In this context, reduced MCC coupling in hearing loss suggests that local temporal organization becomes less closely aligned with the anatomical variation captured by GMV (Del Mauro and Wang, 2025).

A plausible explanation involves the continuing demands that degraded auditory input places on monitoring and adaptive control. Anterior midcingulate territories integrate information relevant to selecting and adjusting behavior under uncertainty (Shackman et al., 2011). In the auditory domain, cingulo-opercular activity increases during difficult word recognition and predicts improved recognition on subsequent trials, supporting a role in adaptive listening control (Vaden et al., 2013). Chronic auditory degradation could therefore alter the balance of inputs and ongoing states influencing cingulate activity, allowing participants with similar regional GMV to exhibit different patterns of spontaneous BOLD recurrence. This interpretation places the local MCC finding within a broader account of experience-dependent functional reorganization, in which temporal activity reflects the accumulated demands of listening as well as regional anatomy. The weak GMV–BEN associations within the right-STG seed further distinguish the two regional findings: STG showed altered BOLD recurrence after local GMV adjustment, whereas MCC showed a change in the relationship between morphology and temporal activity.

The interregional results suggest a corresponding change in the relationship between structural pathways and functional coordination. Using the final QSIPrep tractography estimates, STG–MCC SC–FC coupling was positive in controls and weakly negative in hearing loss, with the exploratory directional comparison supporting stronger coupling in controls. Together with the focal FC increase and the small tractography group effect, this pattern suggests that functional coordination in hearing loss is less closely related to individual differences in the reconstructed pathway. At the network level, A. Wang et al. (2026) reported altered structural network topology and no significant group difference in whole-brain SC–FC coupling in noise-induced hearing loss. Our exploratory STG–MCC analysis focuses on between-person correspondence along a specific auditory–cingulate connection, suggesting that the spatial scale and definition of coupling are important considerations when characterizing hearing-related reorganization. Structural–functional modeling provides a basis for this interpretation: resting-state correlations reflect both direct anatomical connections and interactions mediated through indirect pathways, so functional coupling can change through alterations in distributed network dynamics (Honey et al., 2009). For STG and MCC, candidate mechanisms include more consistent shared input, altered regional responsiveness, and greater involvement of indirect routes linking auditory and control systems. These mechanisms could strengthen temporal covariation while reducing the extent to which between-person FC differences follow tractography-derived connection strength (Honey et al., 2009).

This account is consistent with evidence that hearing loss changes auditory–cingulate connectivity. Fitzhugh et al. (2019) reported stronger resting connectivity between right Heschl’s gyrus and dorsal anterior cingulate cortex with poorer hearing, after adjustment for age and cognitive performance. The present findings extend that auditory–cingulate association by linking a BEN-defined auditory region to a structurally identified MCC region and examining their cross-modal correspondence. Across local and interregional scales, the results suggest that hearing-related reorganization involves a redistribution of the relationship between anatomy and spontaneous function. Auditory cortical activity becomes more temporally regular, while cingulate temporal activity and auditory–cingulate coordination become less closely aligned with the measured structural variation. This perspective shifts attention toward how existing anatomy is functionally engaged under degraded hearing and motivates testing whether these coupling changes track listening effort, speech-in-noise performance, and adaptation to hearing rehabilitation (Fitzhugh et al., 2019).

Taken together, the results suggest that chronic auditory degradation affects both local temporal organization in auditory association cortex and its coordination with a region involved in performance monitoring. More recurrent right-STG activity may reflect repeated engagement of a restricted set of auditory predictions or representations, while the MCC findings may reflect the continuing demand to evaluate and adjust behavior when sensory evidence is unreliable. Neuroimaging studies of challenging speech consistently implicate control systems in maintaining performance, and sustained engagement of these systems contributes to listening effort and fatigue (Eckert et al., 2016). Audiometric severity, speech-in-noise performance, listening effort, and cognition are therefore important variables for determining whether the observed STG-MCC organization supports compensation, reflects inefficiency, or changes across stages of hearing decline.

Several features of the design define the next empirical steps. The cross-sectional data do not establish the temporal sequence between STG and MCC changes. The connectivity and coupling analyses followed corrected findings within the same cohort, and the DCM and coupling evidence remained exploratory. Longitudinal imaging could test whether altered STG temporal organization precedes cingulate recruitment or develops in parallel with it. Replication in another multimodal cohort, together with measures of hearing duration, hearing-aid exposure, tinnitus, speech-in-noise performance, and cognition, would establish the generalizability and clinical meaning of the data-driven auditory-cingulate findings. Directional coupling comparisons were exploratory and selected after inspection of the associations.

## Conclusions

Hearing loss was associated with greater temporal recurrence in right auditory association cortex, accompanied by higher MCC GMV and stronger focal auditory–cingulate functional coordination. Exploratory analyses further suggested weaker local and interregional correspondence between structural variation and spontaneous function. These findings extend the characterization of hearing-related plasticity to the temporal organization of intrinsic activity and its relationship to brain structure. They provide a basis for investigating how altered auditory input changes cortical processing demands and contributes to individual differences in everyday communication.

## Acknowledgements

We thank the investigators and participants of the Hearing Loss Connectome project for making the deidentified multimodal imaging dataset publicly available through OpenNeuro (doi:10.18112/openneuro.ds005026.v1.0.0).

## Ethics statement

The original study was approved by the Institutional Review Board of the University of Salerno. The present study analyzed deidentified data publicly available through OpenNeuro (ds005026).

## Data and code availability

The source imaging dataset is publicly available from OpenNeuro under accession ds005026 (doi:10.18112/openneuro.ds005026.v1.0.0). Analysis code, masks, and derived results will be made available in the accompanying project repository upon publication.

## References

Abraham A, Pedregosa F, Eickenberg M, et al. Machine learning for neuroimaging with scikit-learn. Frontiers in Neuroinformatics. 2014;8:14. doi:10.3389/fninf.2014.00014.

Alfandari D, Vriend C, Heslenfeld DJ, Versfeld NJ, Kramer SE, Zekveld AA. Brain volume differences associated with hearing impairment in adults. Trends in Hearing. 2018;22:2331216518763689.

Andin J, Holmer E. Reorganization of large-scale brain networks in deaf signing adults: The role of auditory cortex in functional reorganization following deafness. Neuropsychologia. 2022;166:108139.

Antonopoulos G, More S, Raimondo F, Eickhoff SB, Hoffstaedter F, Patil KR. A systematic comparison of VBM pipelines and their application to age prediction. NeuroImage. 2023;279:120292. doi:10.1016/j.neuroimage.2023.120292.

Avants BB, Tustison NJ, Song G, Cook PA, Klein A, Gee JC. A reproducible evaluation of ANTs similarity metric performance in brain image registration. NeuroImage. 2011;54:2033–2044. doi:10.1016/j.neuroimage.2010.09.025.

Behrens TEJ, Berg HJ, Jbabdi S, Rushworth MFS, Woolrich MW. Probabilistic diffusion tractography with multiple fibre orientations: What can we gain? NeuroImage. 2007;34:144–155. doi:10.1016/j.neuroimage.2006.09.018.

Behzadi Y, Restom K, Liau J, Liu TT. A component based noise correction method (CompCor) for BOLD and perfusion based fMRI. NeuroImage. 2007;37:90–101. doi:10.1016/j.neuroimage.2007.04.042.

Biswal B, Yetkin FZ, Haughton VM, Hyde JS. Functional connectivity in the motor cortex of resting human brain using echo-planar MRI. Magnetic Resonance in Medicine. 1995;34:537–541. doi:10.1002/mrm.1910340409.

Cieslak M, Cook PA, He X, et al. QSIPrep: an integrative platform for preprocessing and reconstructing diffusion MRI data. Nature Methods. 2021;18:775–778.

Cox RW. AFNI: software for analysis and visualization of functional magnetic resonance neuroimages. Computers and Biomedical Research. 1996;29:162–173. doi:10.1006/cbmr.1996.0014.

Del Mauro G, Wang Z. rsfMRI-based brain entropy is negatively correlated with gray matter volume and surface area. Brain Structure and Function. 2025;230:35. doi:10.1007/s00429-025-02897-6.

Douaud G, Smith S, Jenkinson M, et al. Anatomically related grey and white matter abnormalities in adolescent-onset schizophrenia. Brain. 2007;130:2375–2386. doi:10.1093/brain/awm184.

Eckert MA, Teubner-Rhodes S, Vaden KI Jr. Is listening in noise worth it? The neurobiology of speech recognition in challenging listening conditions. Ear and Hearing. 2016;37 Suppl 1:101S–110S. doi:10.1097/AUD.0000000000000300.

Esteban O, Markiewicz CJ, Blair RW, et al. fMRIPrep: a robust preprocessing pipeline for functional MRI. Nature Methods. 2019;16:111–116. doi:10.1038/s41592-018-0235-4.

Fisher RA. On the probable error of a coefficient of correlation deduced from a small sample. Metron. 1921;1:3–32.

Fitzhugh MC, Hemesath A, Schaefer SY, Baxter LC, Rogalsky C. Functional connectivity of Heschl’s gyrus associated with age-related hearing loss: a resting-state fMRI study. Frontiers in Psychology. 2019;10:2485.

Friston KJ, Kahan J, Biswal B, Razi A. A DCM for resting state fMRI. NeuroImage. 2014;94:396–407.

Garrett DD, Samanez-Larkin GR, MacDonald SWS, Lindenberger U, McIntosh AR, Grady CL. Moment-to-moment brain signal variability: a next frontier in human brain mapping? Neuroscience and Biobehavioral Reviews. 2013;37:610–624. doi:10.1016/j.neubiorev.2013.02.015.

Good CD, Johnsrude IS, Ashburner J, Henson RNA, Friston KJ, Frackowiak RSJ. A voxel-based morphometric study of ageing in 465 normal adult human brains. NeuroImage. 2001;14:21–36. doi:10.1006/nimg.2001.0786.

Gorgolewski K, Burns CD, Madison C, et al. Nipype: a flexible, lightweight and extensible neuroimaging data processing framework in Python. Frontiers in Neuroinformatics. 2011;5:13. doi:10.3389/fninf.2011.00013.

Greve DN, Fischl B. Accurate and robust brain image alignment using boundary-based registration. NeuroImage. 2009;48:63–72. doi:10.1016/j.neuroimage.2009.06.060.

Honey CJ, Sporns O, Cammoun L, et al. Predicting human resting-state functional connectivity from structural connectivity. Proceedings of the National Academy of Sciences of the United States of America. 2009;106:2035–2040. doi:10.1073/pnas.0811168106.

Jenkinson M, Bannister P, Brady M, Smith S. Improved optimization for the robust and accurate linear registration and motion correction of brain images. NeuroImage. 2002;17:825–841. doi:10.1006/nimg.2002.1132.

Jenkinson M, Beckmann CF, Behrens TEJ, Woolrich MW, Smith SM. FSL. NeuroImage. 2012;62:782-790. doi:10.1016/j.neuroimage.2011.09.015.

Lempel A, Ziv J. On the complexity of finite sequences. IEEE Transactions on Information Theory. 1976;22:75–81. doi:10.1109/TIT.1976.1055501.

Li N, Ma W, Ren F, et al. Neurochemical and functional reorganization of the cognitive-ear link underlies cognitive impairment in presbycusis. NeuroImage. 2023;268:119861.

Li X, Fu W, Wang Y, et al. Functional-structural coupling: brain reorganization in presbycusis is related to cognitive impairment. eNeuro. 2026;13(3):ENEURO.0294-25.2026. doi:10.1523/ENEURO.0294-25.2026.

Li Y, Booth JR, Peng D, et al. Altered intra- and inter-regional synchronization of superior temporal cortex in deaf people. Cerebral Cortex. 2013;23:1988–1996.

Lin L, Chang D, Song D, Li Y, Wang Z. Lower resting brain entropy is associated with stronger task activation and deactivation. NeuroImage. 2022;249:118875.

MacKinnon JG, White H. Some heteroskedasticity-consistent covariance matrix estimators with improved finite sample properties. Journal of Econometrics. 1985;29:305–325. doi:10.1016/0304-4076(85)90158-7.

Mesgarani N, Cheung C, Johnson K, Chang EF. Phonetic feature encoding in human superior temporal gyrus. Science. 2014;343:1006–1010. doi:10.1126/science.1245994.

Nichols TE, Holmes AP. Nonparametric permutation tests for functional neuroimaging: a primer with examples. Human Brain Mapping. 2002;15:1–25. doi:10.1002/hbm.1058.

Oganian Y, Chang EF. A speech envelope landmark for syllable encoding in human superior temporal gyrus. Science Advances. 2019;5:eaay6279. doi:10.1126/sciadv.aay6279.

Orepić P, Truccolo W, Halgren E, Cash SS, Giraud AL, Proix T. Top-down semantic predictions align phonetic neuronal dynamics in human superior temporal gyrus. Nature Communications. Published online October 3, 2026. doi:10.1038/s41467-026-77942-x.

Ponticorvo S, Esposito F. Hearing loss Connectome, version 1.0.0 [dataset]. OpenNeuro. 2024. doi:10.18112/openneuro.ds005026.v1.0.0.

Ponticorvo S, Manara R, Cassandro E, et al. Cross-modal connectivity effects in age-related hearing loss. Neurobiology of Aging. 2022;111:1–13.

Richman JS, Moorman JR. Physiological time-series analysis using approximate entropy and sample entropy. American Journal of Physiology Heart and Circulatory Physiology. 2000;278:H2039–H2049. doi:10.1152/ajpheart.2000.278.6.H2039.

Shackman AJ, Salomons TV, Slagter HA, Fox AS, Winter JJ, Davidson RJ. The integration of negative affect, pain and cognitive control in the cingulate cortex. Nature Reviews Neuroscience. 2011;12:154–167. doi:10.1038/nrn2994.

Shang Y, Hinkley LB, Cai C, Mizuiri D, Cheung SW, Nagarajan SS. Cross-modal plasticity in adult single-sided deafness revealed by alpha-band resting-state functional connectivity. NeuroImage. 2020;207:116376.

Shiell MM, Champoux F, Zatorre RJ. Reorganization of auditory cortex in early-deaf people: functional connectivity and relationship to hearing aid use. Journal of Cognitive Neuroscience. 2015;27:150–163.

Sohoglu E, Davis MH. Perceptual learning of degraded speech by minimizing prediction error. Proceedings of the National Academy of Sciences of the United States of America. 2016;113(12):E1747–E1756. doi:10.1073/pnas.1523266113.

Sohoglu E, Peelle JE, Carlyon RP, Davis MH. Predictive top-down integration of prior knowledge during speech perception. Journal of Neuroscience. 2012;32(25):8443–8453. doi:10.1523/JNEUROSCI.5069-11.2012.

Song D, Chang D, Zhang J, Ge Q, Zang YF, Wang Z. Associations of brain entropy (BEN) to cerebral blood flow and fractional amplitude of low-frequency fluctuations in the resting brain. Brain Imaging and Behavior. 2019a;13:1486–1495.

Song D, Chang D, Zhang J, Peng W, Shang Y, Gao X, Wang Z. Reduced brain entropy by repetitive transcranial magnetic stimulation on the left dorsolateral prefrontal cortex in healthy young adults. Brain Imaging and Behavior. 2019b;13:421–429.

Song D, Deng XP, Chang D, Wang Z. Altered resting-state brain entropy by repetitive transcranial magnetic stimulation across the human cortex. Cerebral Cortex. 2025;35(7):bhaf171. doi:10.1093/cercor/bhaf171.

Song D, Wang Z. Age-dependent effects of intranasal oxytocin administration were revealed by resting brain entropy (BEN). Behavioural Brain Research. 2026;500:115985. doi:10.1016/j.bbr.2025.115985.

Su M, Ren F, Li N, et al. Alterations of excitation-inhibition balance and brain network dynamics support sensory deprivation theory in presbycusis. Human Brain Mapping. 2024;45:e70067.

Tong Z, Zhang J, Xing C, et al. Reorganization of the cortical connectome functional gradient in age-related hearing loss. NeuroImage. 2023;284:120475.

Tustison NJ, Avants BB, Cook PA, et al. N4ITK: improved N3 bias correction. IEEE Transactions on Medical Imaging. 2010;29:1310–1320. doi:10.1109/TMI.2010.2046908.

Tzourio-Mazoyer N, Landeau B, Papathanassiou D, et al. Automated anatomical labeling of activations in SPM using a macroscopic anatomical parcellation of the MNI MRI single-subject brain. NeuroImage. 2002;15:273–289. doi:10.1006/nimg.2001.0978.

Vaden KI Jr, Kuchinsky SE, Ahlstrom JB, Dubno JR, Eckert MA. Cortical activity predicts which older adults recognize speech in noise and when. Journal of Neuroscience. 2015;35:3929–3937.

Vaden KI Jr, Kuchinsky SE, Cute SL, Ahlstrom JB, Dubno JR, Eckert MA. The cingulo-opercular network provides word-recognition benefit. Journal of Neuroscience. 2013;33:18979–18986. doi:10.1523/JNEUROSCI.1417-13.2013.

Vogt BA. Midcingulate cortex: structure, connections, homologies, functions and diseases. Journal of Chemical Neuroanatomy. 2016;74:28–46.

Wang A, Bao X, Wang C, Huang R, Wang L, Lv M, Zhang G. Alterations of brain network topology and structural-functional connectivity coupling in noise-induced hearing loss based on macroscopic scale. Frontiers in Neuroscience. 2026;19:1661096. doi:10.3389/fnins.2025.1661096.

Wang X, Ao B, Yazdani Y, Lau C, Abouzari M, Ratnanather JT, Manno FAM. Meta-analysis and meta-regression of hearing loss fMRI activity, connectivity, and fluctuation alterations: Heterogeneity of auditory sensory impact across the lifespan. NeuroImage. 2026;336:121973. doi:10.1016/j.neuroimage.2026.121973.

Wang X, Xu P, Li P, et al. Alterations in gray matter volume due to unilateral hearing loss. Scientific Reports. 2016;6:25811.

Wang Z, Li Y, Childress AR, Detre JA. Brain entropy mapping using fMRI. PLoS ONE. 2014;9:e89948.

Watanabe T, Rees G, Masuda N. Atypical intrinsic neural timescale in autism. eLife. 2019;8:e42256. doi:10.7554/eLife.42256.

Winkler AM, Ridgway GR, Webster MA, Smith SM, Nichols TE. Permutation inference for the general linear model. NeuroImage. 2014;92:381–397. doi:10.1016/j.neuroimage.2014.01.060.

Wolff A, Berberian N, Golesorkhi M, Gomez-Pilar J, Zilio F, Northoff G. Intrinsic neural timescales: temporal integration and segregation. Trends in Cognitive Sciences. 2022;26:159–173. doi:10.1016/j.tics.2021.11.007.

Yang M, Chen HJ, Liu B, et al. Brain structural and functional alterations in patients with unilateral hearing loss. Hearing Research. 2014;316:37–43.

Zang YF, He Y, Zhu CZ, et al. Altered baseline brain activity in children with ADHD revealed by resting-state functional MRI. Brain and Development. 2007;29:83–91. doi:10.1016/j.braindev.2006.07.002.

Zang YF, Jiang TZ, Lu YL, He Y, Tian LX. Regional homogeneity approach to fMRI data analysis. NeuroImage. 2004;22:394–400. doi:10.1016/j.neuroimage.2003.12.030.

Zeidman P, Jafarian A, Seghier ML, et al. A guide to group effective connectivity analysis, part 2: Second level analysis with PEB. NeuroImage. 2019;200:12–25. doi:10.1016/j.neuroimage.2019.06.032.

Zhang Y, Brady M, Smith S. Segmentation of brain MR images through a hidden Markov random field model and the expectation-maximization algorithm. IEEE Transactions on Medical Imaging. 2001;20:45–57. doi:10.1109/42.906424.

Zou QH, Zhu CZ, Yang Y, et al. An improved approach to detection of amplitude of low-frequency fluctuation for resting-state fMRI: fractional ALFF. Journal of Neuroscience Methods. 2008;172:137–141. doi:10.1016/j.jneumeth.2008.04.012.

