## Supplementary Materials for "Reduced Resting Brain Entropy and Structural-Functional Reorganization in Hearing Loss"

### Supplementary Methods

#### Complementary resting-state measures

The first four BOLD volumes were removed, leaving 396 observations. ALFF and fALFF used the 0.009-0.08 Hz band; the fALFF denominator contained nonzero frequencies from 0 to 0.25 Hz. ReHo was calculated as Kendall concordance across each voxel and its 26 neighbors. Lempel-Ziv complexity used median-binarized time series and normalized LZ76 complexity. MSSD quantified successive-sample variability, and intrinsic timescale was calculated from the positive autocorrelation area to the first zero crossing, with a maximum lag of 50 s. All maps were smoothed at 8 mm FWHM.

Within left and right AAL STG masks, voxelwise models included age, sex, mean framewise displacement, TIV, and voxelwise 8-mm-FWHM GMV. Clusters were formed at two-sided p<0.001 and tested with 5,000-permutation maximum-cluster FWER correction separately in each hemisphere. For whole-brain regional analyses, ALFF, fALFF, ReHo, MSSD, and intrinsic timescale were averaged within 116 AAL regions. Group models included age, sex, mean framewise displacement, TIV, and regional GMV. A 10,000-permutation max-|t| procedure controlled FWER jointly across 580 tests. Metric-wise FDR analyses were examined as sensitivity tests.

#### Structure-function coupling

Local GMV-BEN partial correlations controlled age, sex, and mean FD. SC-FC correlations used final QSIPrep normalized tract values within the 28-voxel exploratory MCC target, log-transformed as ln(1 + 10^6 × SC), and additionally controlled TIV and QSIPrep seed voxel count. Within-group tests used n-k-2 degrees of freedom. Exploratory directional Fisher tests compared NH > HL correlations using variance 1/(n-k-3) for each group. The direction was selected after inspection of the associations. Coupling p values were uncorrected. Balanced subsampling retained all controls and repeatedly sampled 30 hearing-loss participants without replacement; percentile ranges summarize sensitivity to downsampling.

#### Supportive localization and robustness

The supportive GMV voxelwise analysis used the same 8-mm GMV maps with age, sex, and TIV adjustment, two-sided voxelwise p < 0.001, 26-neighbor connectivity, and 5,000-permutation maximum-cluster-extent FWER correction within bilateral MCC. BEN leave-one-out selection refitted the voxelwise model including local GMV in each training set; clusters required at least 10 voxels. Held-out BEN and GMV were available in 79 of 82 folds. Figure 2A displays voxel-selection frequency over all 82 folds. For FC, the five most positive MCC voxels were selected in each training set while the STG seed remained fixed. Head-motion sensitivity added FD squared or excluded participants in the upper 5% or 10% of mean FD.

### Supplementary Results

#### STG specificity analyses

None of the six complementary measures produced a corrected cluster in either STG with 8-mm voxelwise GMV adjustment. The uncorrected candidates included ALFF (R STG, 6 voxels; peak t = 3.790, cluster FWER p = 0.060); Lempel-Ziv complexity (L STG, 2 voxels; peak t = 3.700, cluster FWER p = 0.139); intrinsic timescale (L STG, 1 voxel; peak t = 3.429, cluster FWER p = 0.105).

**Supplementary Table S1.** Complementary resting-state measures in the STG. HL, hearing loss; NH, normal hearing.

| **Metric** | **STG result** | **Direction** | **Peak t** | **Corrected result** |
| --- | --- | --- | --- | --- |
| ALFF | R STG, 6 voxels | HL > NH | 3.790 | cluster FWER p=0.060 |
| fALFF | No p<0.001 cluster | - | - | Not significant |
| ReHo | No p<0.001 cluster | - | - | Not significant |
| Lempel-Ziv | L STG, 2 voxels | HL > NH | 3.700 | cluster FWER p=0.139 |
| MSSD | No p<0.001 cluster | - | - | Not significant |
| Intrinsic timescale | L STG, 1 voxel | HL > NH | 3.429 | cluster FWER p=0.105 |

#### Whole-brain regional analyses

No regional effect survived joint maxT-FWER correction across five metrics and 116 AAL regions. The smallest corrected p value was observed for lower intrinsic timescale in left posterior cingulate cortex. The strongest result for each metric is shown below; the complete 580-test table accompanies this document as Supplementary Data S1.

**Supplementary Table S2.** Strongest AAL regional effect for each complementary metric under the joint 580-test correction.

| **Metric** | **AAL region** | **Direction** | **t** | **p unc** | **p FWE** |
| --- | --- | --- | --- | --- | --- |
| ALFF | Temporal_Pole_Mid_R | HL > NH | 3.134 | 0.002 | 0.517 |
| fALFF | Frontal_Inf_Orb_R | HL > NH | 3.136 | 0.002 | 0.516 |
| ReHo | Cerebelum_8_L | HL > NH | 3.832 | <0.001 | 0.095 |
| MSSD | Cingulum_Post_L | HL > NH | 3.812 | <0.001 | 0.101 |
| Intrinsic timescale | Cingulum_Post_L | HL < NH | -3.850 | <0.001 | 0.089 |

Metric-wise FDR sensitivity analyses identified 10 regional effects after 8-mm GMV adjustment, predominantly involving cerebellar ReHo. The complete regional estimates are reported in Supplementary Table S3. These effects did not survive the joint five-metric correction.

**Supplementary Table S3.** Metric-wise ReHo FDR sensitivity results after regional GMV adjustment.

| **Region** | **Direction** | **beta** | **t** | **p unc** | **q FDR** |
| --- | --- | --- | --- | --- | --- |
| Cerebelum_8_L | HL > NH | 0.226 | 3.832 | <0.001 | 0.028 |
| Cerebelum_8_R | HL > NH | 0.193 | 3.606 | <0.001 | 0.028 |
| Frontal_Sup_R | HL < NH | -0.156 | -3.522 | <0.001 | 0.028 |
| Frontal_Mid_R | HL < NH | -0.161 | -3.331 | 0.001 | 0.039 |
| Vermis_9 | HL > NH | 0.184 | 3.164 | 0.002 | 0.041 |
| Cerebelum_10_R | HL > NH | 0.163 | 3.156 | 0.002 | 0.041 |
| Cerebelum_10_L | HL > NH | 0.186 | 3.083 | 0.003 | 0.041 |
| Cerebelum_9_L | HL > NH | 0.220 | 3.065 | 0.003 | 0.041 |
| Cerebelum_3_L | HL > NH | 0.207 | 3.015 | 0.004 | 0.041 |
| Cerebelum_9_R | HL > NH | 0.191 | 3.013 | 0.004 | 0.041 |

#### Coupling statistics

Supplementary Table S4. Group-specific partial correlations and exploratory directional Fisher comparisons. NH n=30; HL n=52. Local models removed three covariates; interregional models removed five covariates.

| Region / pair | NH r | NH df | NH p | HL r | HL df | HL p | NH > HL p |
| --- | --- | --- | --- | --- | --- | --- | --- |
| Right STG GMV-BEN | 0.009 | 25 | 0.965 | 0.115 | 47 | 0.430 | 0.664 |
| Right MCC GMV-BEN | 0.418 | 25 | 0.030 | -0.014 | 47 | 0.924 | 0.034 |
| Bilateral MCC GMV-BEN | 0.388 | 25 | 0.046 | -0.057 | 47 | 0.697 | 0.032 |
| STG-MCC SC-FC | 0.377 | 23 | 0.063 | -0.070 | 45 | 0.642 | 0.037 |

#### Motion sensitivity and held-out estimates

Supplementary Table S5. Sensitivity of BEN and FC summary effects to head motion and held-out ROI selection. BEN models include local GMV; FC models use the focal five-voxel target.

| Outcome | Analysis | n | t | df | p |
| --- | --- | --- | --- | --- | --- |
| BEN | primary | 82 | -4.299 | 75 | <0.001 |
| BEN | plus FD squared | 82 | -4.304 | 74 | <0.001 |
| BEN | exclude top 5% FD | 77 | -4.472 | 70 | <0.001 |
| BEN | exclude top 10% FD | 73 | -4.347 | 66 | <0.001 |
| FC | primary | 82 | 4.395 | 76 | <0.001 |
| FC | plus FD squared | 82 | 4.362 | 75 | <0.001 |
| FC | exclude top 5% FD | 77 | 3.742 | 71 | <0.001 |
| FC | exclude top 10% FD | 73 | 3.414 | 67 | 0.001 |
| BEN LOO | held out | 79 | -2.837 | 72 | 0.006 |
| FC LOO | held out | 82 | 3.560 | 76 | <0.001 |

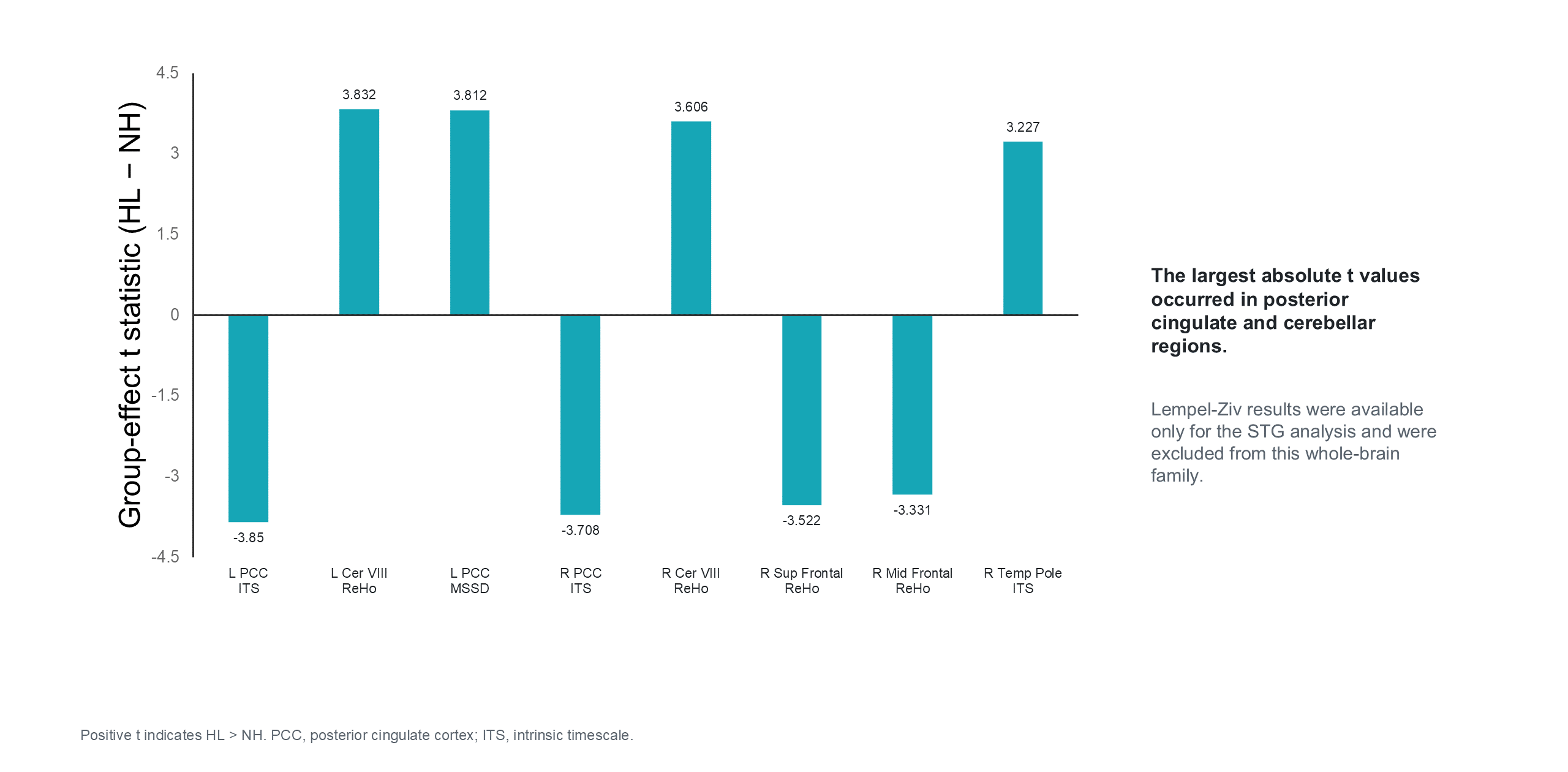

Supplementary Figure S1. Whole-brain AAL sensitivity analysis of auxiliary resting-state metrics. The eight largest absolute group-effect t statistics across the five measures and 116 AAL regions are displayed. These involved intrinsic timescale, ReHo, and mean squared successive difference in posterior cingulate, cerebellar, frontal, and temporal-pole regions. The smallest joint maximum-statistic FWER-corrected p value was 0.089 across five metrics and 116 AAL regions (580 tests). Lempel-Ziv measures were available only for the STG analysis and were analyzed separately from this whole-brain correction family.

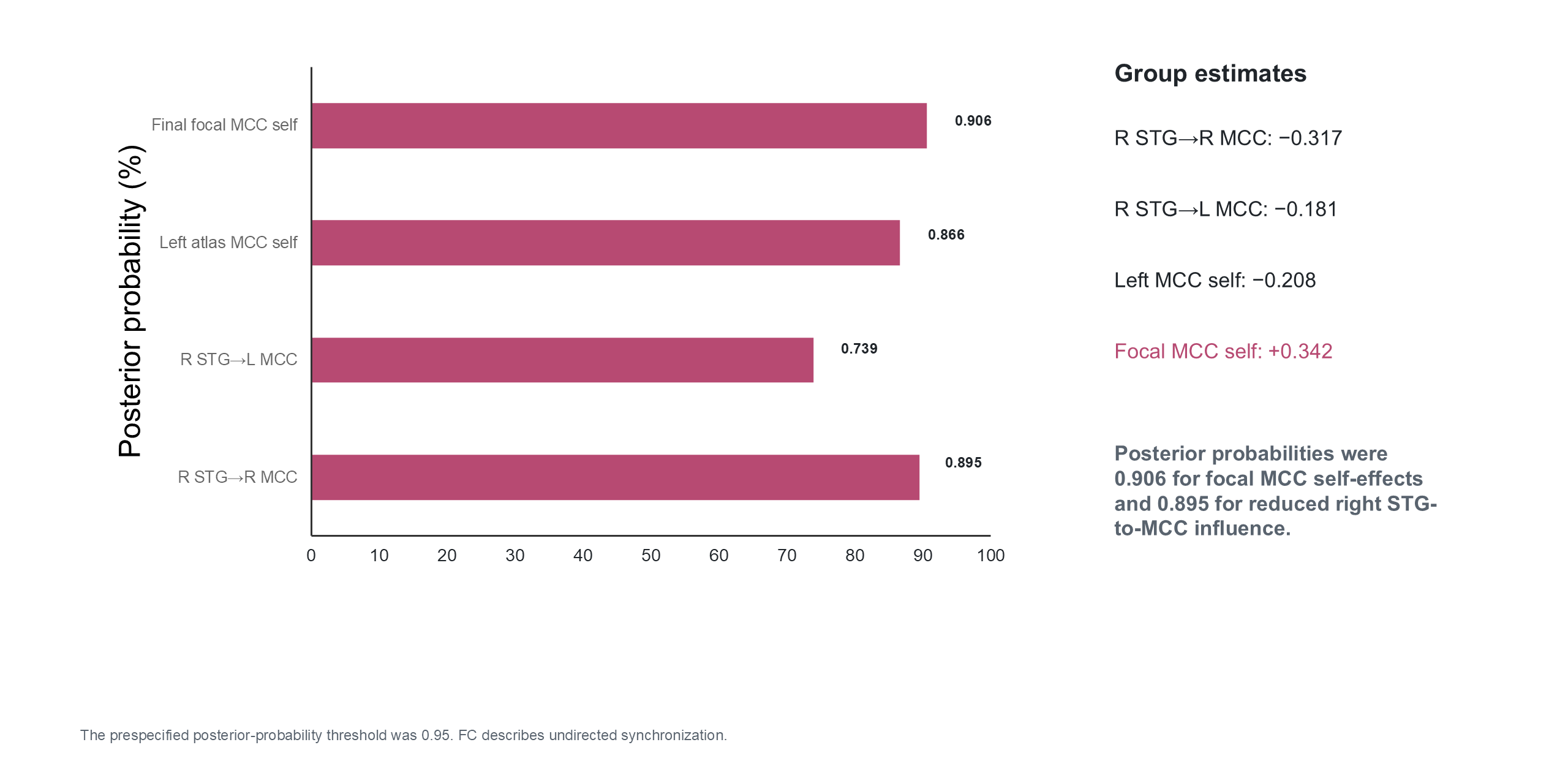

Supplementary Figure S2. Spectral dynamic causal modeling of STG-MCC interactions. Bayesian model averages summarize hearing-loss versus normal-hearing effects for the separate right-STG–left-MCC and right-STG–right-MCC atlas-based models and the final right-STG–focal-MCC model. The largest posterior evidence involved focal-MCC self-effects (estimate=0.342, posterior probability=0.906) and reduced right-STG-to-MCC influence (estimate=-0.317, posterior probability=0.895); the posterior-probability criterion was 0.95. Displayed estimates are from the sensitivity models retaining participants with explained spectral variance ≥70%.

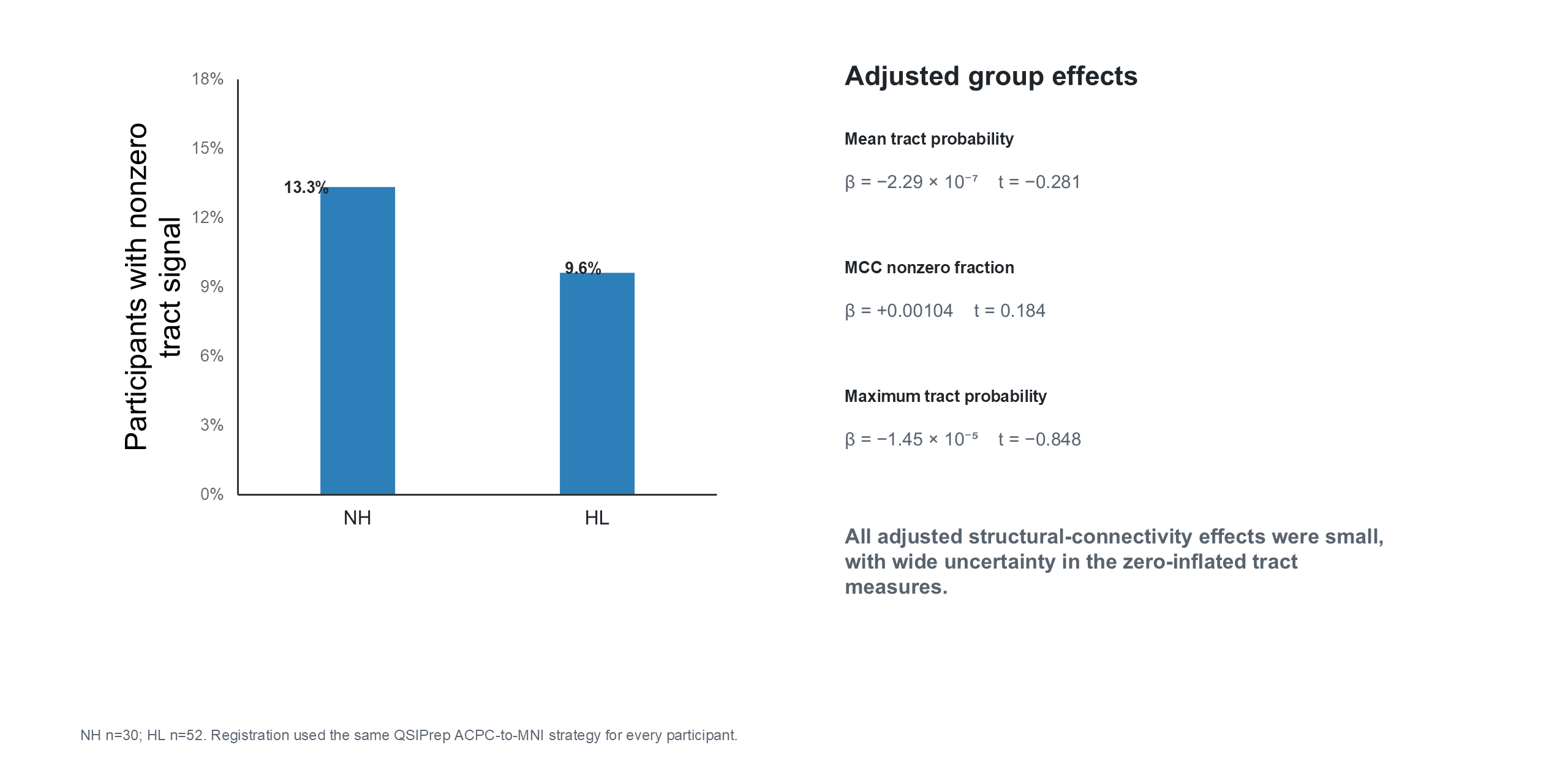

Supplementary Figure S3. QSIPrep-based right STG-MCC probabilistic tractography. Participant-level tract measures were highly zero-inflated. Models included age, sex, and the subject-specific QSIPrep STG seed voxel count. Adjusted effects were small for mean tract probability (beta=-2.29 x 10^-7, p=0.780), MCC nonzero fraction (beta=0.00104, p=0.855), and maximum tract probability (beta=-1.45 x 10^-5, p=0.399), with wide uncertainty across the zero-inflated measures.

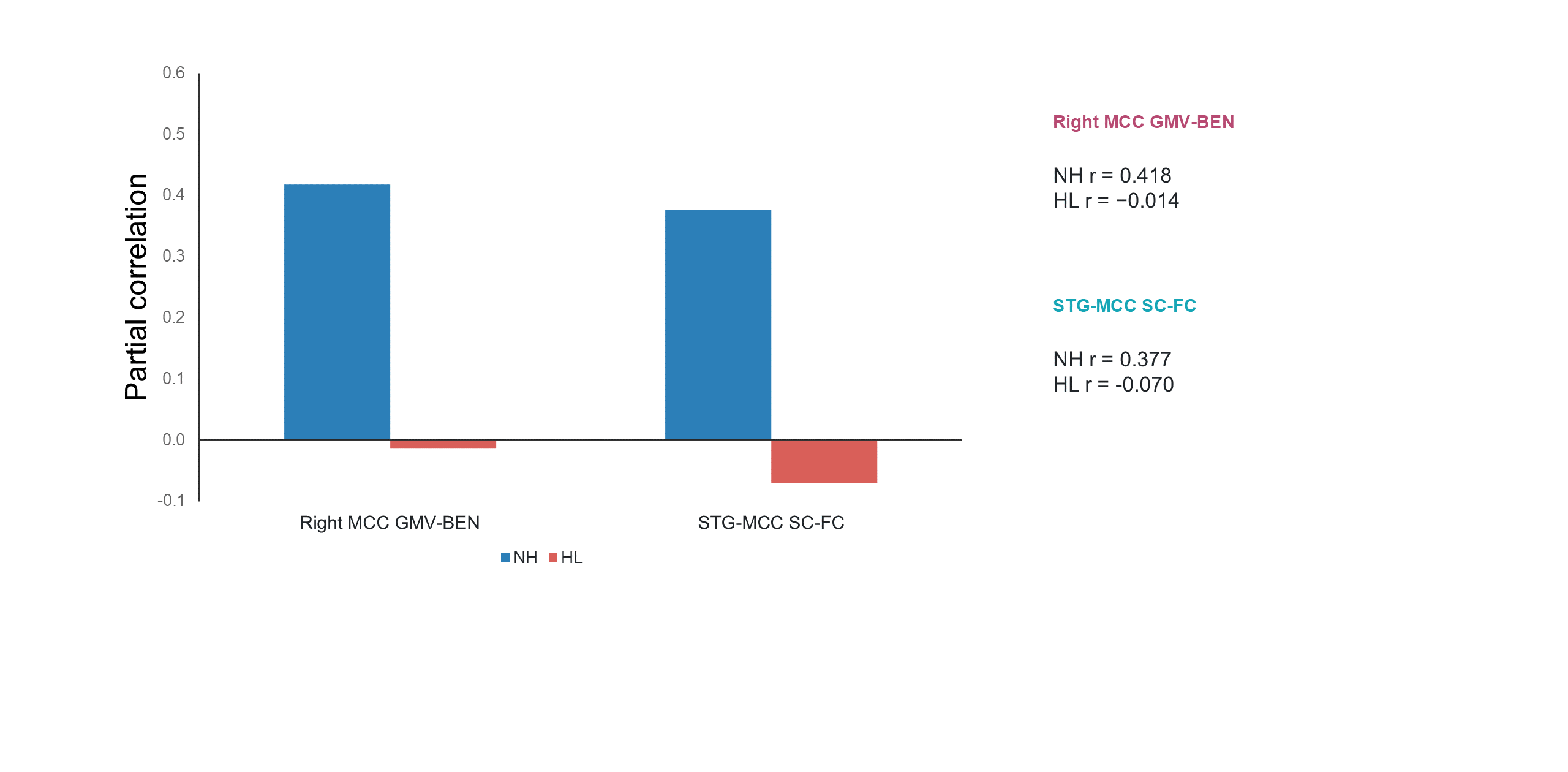

Supplementary Figure S4. Group-specific local and interregional coupling. Partial correlations summarize right-MCC GMV-BEN coupling and STG-MCC structural-functional coupling within each group. For right-MCC GMV-BEN coupling from independently generated 8-mm maps, NH r=0.418, p=0.030 and HL r=-0.014, p=0.924; the directional NH > HL Fisher comparison yielded z=1.824 and one-sided p=0.034. For final QSIPrep-based STG-MCC SC-FC coupling, NH r=0.377, p=0.063 and HL r=-0.070, p=0.642; the directional NH > HL Fisher comparison yielded z=1.787 and one-sided p=0.037.
